# Enhancing Simulations With Intra-Subject Variability for Improved Psychophysical Assessments

**DOI:** 10.1101/285387

**Authors:** Mike D. Rinderknecht, Olivier Lambercy, Roger Gassert

## Abstract

Psychometric properties of perceptual assessments, like reliability, depend on the stochastic properties of psychophysical sampling procedures resulting in method variability, as well as inter- and intra-subject variability. Method variability is commonly minimized by optimizing sampling procedures through computer simulations. Inter-subject variability is inherent in the population of interest and cannot be acted upon. In contrast, intra-subject variability introduced by confounds cannot be simply quantified from experimental data, as this data also includes method variability. Therefore, this aspect is generally neglected when developing assessments. Yet, comparing method variability and intra-subject variability could give insights on whether effort should be invested in optimizing the sampling procedure, or in addressing potential confounds instead. We propose a new approach to estimate and model intra-subject variability of psychometric functions by combining computer simulations and behavioral data, and to account for it when simulating experiments. The approach was illustrated in a real-world scenario of proprioceptive difference threshold assessments. The behavioral study revealed a test-retest reliability of 0.212. Computer simulations lacking intra-subject variability predicted a reliability of 0.768, whereas the new approach including an intra-subject variability model lead to a realistic estimate of reliability (0.207). Such a model also allows computing the theoretically maximally attainable reliability (0.552) assuming an ideal sampling procedure. Comparing the reliability estimates when exclusively accounting for method variability versus intra-subject variability reveals that intra-subject variability should be reduced by addressing confounds and that only optimizing the sampling procedure may be insufficient to achieve a high reliability. The new approach also allows accelerating the development of assessments by simulating the converging behavior of the reliability confidence interval with a large number of subjects and retests without requiring additional experiments. Having such a tool of predictive value is especially important for target populations where time is scarce, such as for assessments in clinical settings.

## 1 Introduction

The development of assessments of human perception thresholds (e.g., visual, auditory, tactile, or proprioceptive stimuli) is a challenging field, as these require good psychometric and clinimetric properties such as high reliability, for both research and clinical applications. The selection and optimization of psychophysical assessments is, in general, a lengthy, iterative, and cumbersome process where different psychophysical methods need to be tested and their parameters tuned (Rinderknecht et al., in preparation). Evaluating such procedures requires time and financial resources, as it involves repeated assessment of a large number of subjects. This may present a serious hurdle for the development of reliable assessments, especially for sample populations where available time is scarce and recruitment is difficult or expensive (e.g., neurological patients).

When evaluating and optimizing psychophysical methods (e.g., for a high test-retest reliability), different factors play an essential role: method variability as well as inter- and intra-subject variability. While inter-subject variability clearly has an effect on reliability (Streiner and Norman, 2008), it is given by the population of interest and cannot be acted upon. Previous works have suggested that a lack of correlation, respectively agreement, between methods tested on the same subjects may originate either from inherent method variability (i.e., based on the stochastic process, the statistical properties of the method, and number of trials) or from intra-subject variability (Rinderknecht et al., 2014). As both method and intra-subject variability are confounded in the outcome measure of a perception assessment, it is impossible to discern one factor from the other and quantify them independently through behavioral studies.

The detection or discrimination capability of physical stimuli is often assumed to resemble a sigmoidal psychometric function (Gescheider, 1997; Macmillan and Douglas Creelman, 2005) based on which the subject generates responses in a stochastic process. Therefore, perception and psychophysical procedures (i.e., complete perception experiments) can be modeled. As a matter of fact, the method variability as well as other performance metrics such as bias and efficiency can be quantified using computer simulations and have been widely investigated for various procedures (Taylor and Douglas Creelman, 1967; Taylor, 1971; Findlay, 1978; Pentland, 1980; Hall, 1981; Madigan and Williams, 1987; Simpson, 1989; Watson and Fitzhugh, 1990; Kaernbach, 1991; Green, 1993; King-Smith et al., 1994)(Rinderknecht et al., in preparation).

In contrast, intra-subject variability is difficult to estimate and cannot be directly quantified based on experimental or simulated data only. Therefore, intra-subject variability has received little attention so far, and is generally neglected in computer simulations. As a consequence, simulations of psychophysical experiments are hardly realistic, and results are not representative. Better knowledge about human perception and the ability to model the intra-subject variability would offer many possibilities, such as comparing, selecting, and tuning different psychophysical methods in simulated scenarios corresponding closely to the real application and population of interest. Furthermore, model-based extrapolation to a larger number of trials or increased sample size, for example to explore their impact on reliability, could then be performed purely in simulation. This could significantly speed up the development and testing of psychophysical assessment procedures.

The aim of this work is twofold: firstly, to present an approach to quantify intra-subject variability, and secondly, to apply and illustrate the approach by creating a general model of intra-subject variability—in this case of proprioceptive perception at the wrist assessed in a two-alternative forced-choice (2AFC) setting. To estimate the intra-subject variability for different parameters of the psychometric function, a dataset with repeated measures from a behavioral study is required. Based on this experimental data, models of the subject’s psychometric function are created to simulate the same population. We propose to add individual, statistical noise models on the different parameters (threshold and slope) of the psychometric functions to simulate intra-subject variability. The level of intra-subject variability (i.e., noise) on the different parameters can be quantified by matching the test-retest reliability of the simulated experiment with the testretest reliability of the behavioral data and by maximizing the similarity between the distributions of outcome measures.

## 2 Materials and methods

### 2.1 Behavioral data

#### 2.1.1 Subjects

Thirty-three healthy young subjects (*N*_subjects_ = 33) were recruited and participated in this experiment (age mean ± SD: 24.1 ± 3.4 years, 20 male and 13 female, 27 right handed, 5 left handed, and 1 ambidextrous). Handedness was assessed with the Edinburgh Handedness Inventory (Oldfield, 1971). Exclusion criteria comprised sensory and motor deficits affecting normal wrist and hand function, as well as any history of neurological or wrist injury. Prior to participating in the experiment, all subjects gave written informed consent. The study was approved by the institutional ethics committee of the ETH Zurich (EK 2015-N-03).

#### 2.1.2 Protocol of the proprioceptive assessment

Each trial of the assessment aiming at estimating the difference threshold of the angular position at the right wrist joint consisted of the consecutive presentation of two different angles and the subsequent judgment by the subject which of the two presented movements was larger (two-interval 2AFC paradigm (Macmillan and Douglas Creelman, 2005)). The movements were applied to the passive wrist with a one degree-of-freedom robotic wrist interface (Chapuis et al., 2010). The movements always started from the resting position (hand aligned with forearm, 0°) and went into flexion direction (maximum 40°). The two presented angles were always centered around a reference of 20°. The difference between the two angles (referred to as level) was defined by an adaptive sampling procedure named Parameter Estimation by Sequential Testing (PEST) (Taylor and Douglas Creelman, 1967). PEST was used with a logarithmic adaptation for positive-only stimuli to avoid an undesired behavior of the algorithm due to zero crossings (Rinderknecht et al., 2014). This adaptive algorithm takes the judgments (also referred to as responses) of past trials into account and changes the difference between the angles accordingly, using heuristic rules to approach the difference threshold as rapidly as possible. The same proprioceptive assessment has been previously used and described in more detail in other studies with a different robotic device for the assessment of the metacarpophalangeal joint (Rinderknecht et al., 2014, 2017, 2018)(Rinderknecht et al., submitted). The same movement timing characteristics and parameters for the PEST algorithm were used in the present experiment, except for the maximum number of trials (start level *x*_0_ = 5.5°, start step Δ*x*_0_ = ±2°, target performance *P*_*t*_ = 75%, Wald sequential likelihood ratio test parameter *W* = 1, minimum step Δ*x*_min_ = ±0.1°, maximum trials at same level *trials*_max@*x*_ = 20, maximum trials in total *trials*_max_ = 120). To avoid any visual or auditory cues (e.g., noise emitted by the motor), the tested arm was occluded from vision and white noise was played over headphones during the whole experiment. Each subject performed the assessment in five session on different days (from 1 to 4 days between sessions, with a maximum of 7 days from the first to the last session).

#### 2.1.3 Estimation of the psychometric function

Based on the data from the assessment sequence (i.e., difference between the two presented angles and corresponding response of the subject), the proportion of correct responses can be calculated for the different levels *x* to fit a sigmoidal psychometric function *ψ*(*x*) (**Figure 1**) using a Maximum Likelihood criterion implemented in the Palamedes MATLAB routines (Prins and Kingdom, 2009):

**Figure 1.**
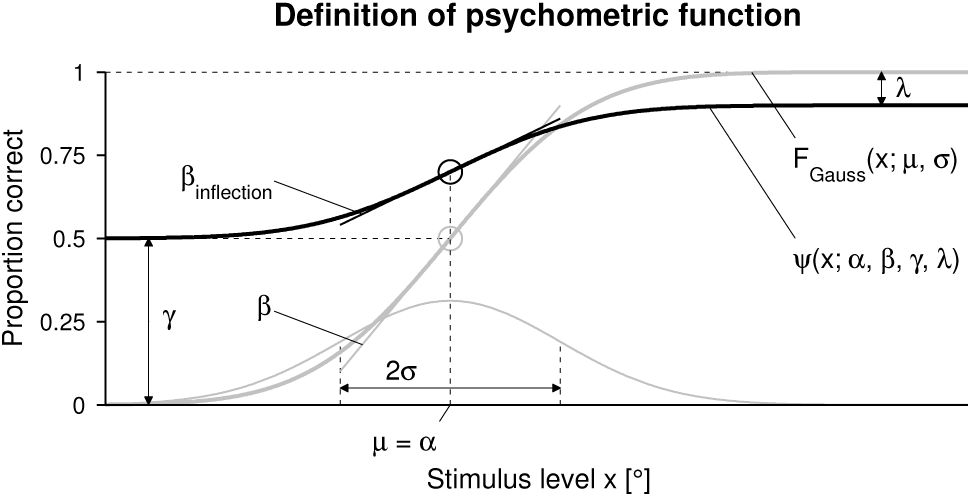
Definition of psychometric function and its parameters. Psychometric function *ψ*(*x*; *α, β, γ, λ*) (bold black sigmoid) and cumulative Gaussian function *F*_Gauss_(*x*; *µ, σ*) (bold gray sigmoid) in the case of a two-alternative forced choice (2AFC) task. The thin gray curve is the underlying Gaussian probability density function. The inflection points are indicated as circles in the respective colors.

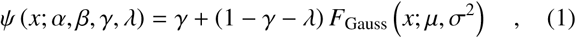

with *F*_Gauss_(*x*; *µ, σ*^2^) a sigmoidal cumulative Gaussian function. In the present work, the threshold parameter *α* corresponds to the mean *µ* of the underlying Gaussian function, and the slope parameter *β* is inversely proportional to the standard deviation *σ*:

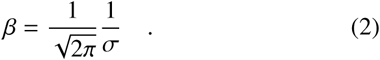

The guess rate parameter *γ* was fixed to 0.5 (according to the 2AFC paradigm), and the lapse rate parameter *λ* was allowed to vary ∈ [0, 0.1]. This has been shown to reduce estimation bias introduced by isolated scattered lapses (Wichmann and Hill, 2001). The actual slope (first order derivative) of *ψ*(*x*) at the inflexion point *α* Is

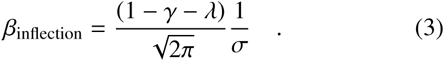

This definition of the slope carries as units one over the units of the stimulus, in the present work [1*/*°], and can be used to compare the slope values across studies using different types of sigmoidal functions *F*(*x*) (Strasburger, 2001). To do arithmetic calculations on the slope (e.g., arithmetic mean), it is reasonable to normalize the slope with the following nonlinear function to a range [0, 1] with arbitrary units [a.u.]:

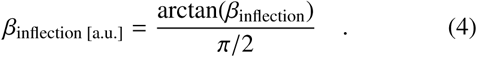

If this nonlinear transformation is not performed, errors in slope estimation can diverge towards infinite for two almost identically steep psychometric functions, which would lead to a distortion when comparing to errors in shallow psychometric functions.

Based on computer simulations, the estimation quality of psychophysical sampling procedures can be calculated for the parameters of the estimated psychometric function depending on the real parameters (Rinderknecht et al., in preparation). Following this work, the estimation performance of PEST was evaluated with computer simulations using the same parameter values as used in the present behavioral study. The variable estimation error cannot be corrected for. However, the average bias (i.e., constant estimation error) can be removed after fitting the psychometric function with the Maximum Likelihood criterion. While PEST can be considered a bias-free sampling procedure for the threshold estimates, the slope estimation bias showed a strong dependence on the real slope and was approximatively corrected by using the following equation:

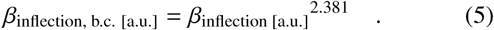

A further estimation bias in psychophysical experiments with human subjects can arise from longer inattention periods, as loss of attention may alter perception (Leek et al., 1991; Fründ et al., 2011; Cohen and Maunsell, 2011). A method to detect and remove such inattention periods in PEST sequences has recently been proposed (Rinderknecht et al., 2018). This method has shown to reduce estimation errors by up to around 75% and was applied *post-hoc* on the PEST sequences recorded in the behavioral study before fitting the psychometric function.

### 2.2 Computer simulations

#### 2.2.3 Population model and templates

A model of the population distribution was created for each parameter of the psychometric function based on the averaged parameters (across measurements) of the psychometric functions obtained in the behavioral study. For the threshold *α* and lapse rate *λ*, the arithmetic mean was calculated for each subject (across the five measurements) to obtain an improved estimate of the subject’s real psychometric function. The same was done for the slope *β*, however, *β* was first converted to the slope at inflection *β*_inflection_ (with the five corresponding lapse rates of the individual subject), normalized (*β*_inflection [a.u.]_), and the bias was removed (*β*_inflection, b.c. [a.u.]_) before averaging across the five measurements. Subsequently, the slope was converted back with the inverse transformations (with the averaged lapse of the individual subject). Averaging was not necessary for the guess rate *γ*, as it was always fixed to 0.5. From these empirical parameter distributions a set of simulated perception models (also referred to as templates *ψ*(*x*)*^T^*) was randomly sampled. To differentiate between psychometric functions and their parameters originating from the behavioral study and the simulated psychometric functions, the symbol *T* was added for variables referring to simulated templates (e.g., *α*^*T*^). The number of templates *ψ*^*T*^ (*x*) was set to be identical to the number of assessed subjects in the behavioral study (*N*_templates_ = 33).

#### 2.2.2 Noise model

The threshold including intra-subject variability was modeled with a log–normal distribution with a support [0, +∞):

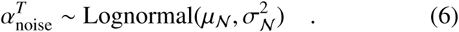

To avoid bias when introducing noise, the mean *µ*_noise_ was defined to be the threshold of the template:

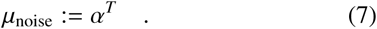

The standard deviation *σ*_noise_ of the variability was controlled with the parameter *ν*_*α*_ ∈ [0, +∞):

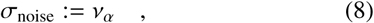

The two parameters of the log–normal distribution were calculated using *µ*_noise_ and the desired *σ*_noise_:

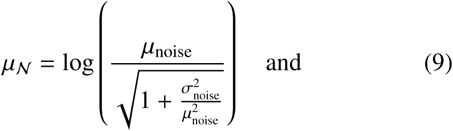

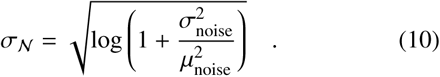

The slope including intra-subject variability was modeled with a beta distribution with a support [0, 1]:

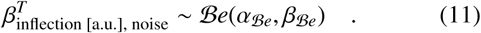

The mean *µ*_noise_ of ℬ*e*(*α*_ℬ*e*_, *β*_ℬ*e*_) was defined to correspond to the normalized slope at the inflection of the template:

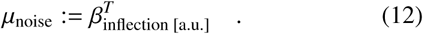

The standard deviation *σ*_noise_ of the variability was controlled with the parameter *ν*_*β*_ ∈ (0, 1] serving as a scaling parameter:

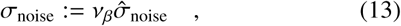

where 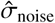 is the maximum possible value for *σ*_noise_ to avoid a U-shaped distribution. This can be guaranteed with at least one of the parameters *α*_ℬ*e*_ or *β*_ℬ*e*_ ≥ 1, leading to:

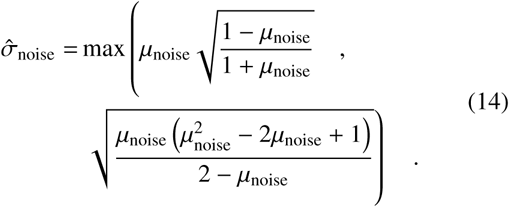

With *µ*_noise_ and *σ*_noise_, the two parameters of the beta distribution ℬ*e*(*α*_ℬ*e*_, *β*_ℬ*e*_) could be calculated:

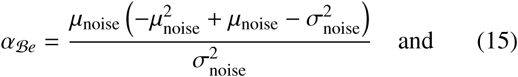

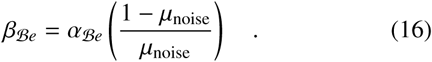

No noise was modeled on the lapse rate *λ*^*T*^ and on the guess rate *γ*^*T*^ = 0.5. The psychometric functions to be used for the simulated PEST sequences were of the form 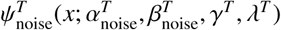 For the threshold, 16 equally distributed noise levels *ν*_*α*_ ∈ [0, 1.5], and for the slope, 14 noise levels *ν*_*β*_ ∈ [0, 1] with a twice as high grid density ∈ [0.7, 1], were simulated.

#### 2.2.3 Procedure

For each combination of *ν*_*α*_ and *ν*_*β*_, the PEST sequence of the 2AFC task was simulated five times for the whole set of templates 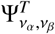 For each single simulated sequence, new random variables 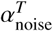 and 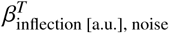 were drawn from the log–normal and beta distributions, respectively, simulating intra-subject variability across the five measurements. The identical PEST parameters as in the behavioral study were used for the computer simulations. Responses to a specific level *x* were simulated by comparing a randomly generated number ∈ 𝒰 (0, 1) to 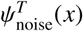 of the respective template. A smaller random number generated a correct response, and a larger random number a false response.

The simulation of the whole set 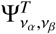 was repeated for each combination of *ν*_*α*_ and *ν*_*β*_ *N*_simulations_ = 1000 times with new randomly sampled parameters (i.e., *α*^*T*^, *β*^*T*^, *γ*^*T*^, *λ*^*T*^) from the population distribution models.

The psychometric functions from the simulated PEST sequences were estimated identically to the behavioral study, including the bias correction. The only difference lay in the inattention correction algorithm (Rinderknecht et al., 2018), which was not applied on the simulated data. It was assumed that significant biases from potential inattention periods in the behavioral study were already corrected for before creating the population model for the templates. Thus, as no inattention periods were modeled in the simulations, there was no need to apply the algorithm. The computer simulations and estimations of the psychometric function were performed

### 2.3 Data analysis

Test-retest reliability of the estimated thresholds from the five measurements of the behavioral study was quantified by computing the ICC(2,1) intraclass correlation coefficient *r* (two-way layout with random effects for absolute agreement) (Shrout and Fleiss, 1979) and its 95% confidence interval (CI) (Lexell and Downham, 2005; de Vet et al., 2006).

Identically, for each set 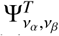, distributions of *N*_simulations_ values for the reliability of the estimated thresholds as well as its lower and upper CI bounds for each combination of *ν*_*α*_ and *ν*_*β*_ were generated. From these *N*_simulations_ reliability values, the arithmetic mean 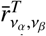 was calculated. In this two-dimensional noise space an iso-reliability contour where the reliability of the simulated experiment matched the reliability of the behavioral study 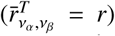 was calculated (set of *ν*_*α*_ and *ν*_*β*_ pairs). To obtain a smoother contour, the reliability surface was interpolated with a spline on a finer grid (by halving the grid intervals three times in each dimension).

To find which *ν*_*α*_ and *ν*_*β*_ pair of the iso-reliability contour corresponds the best to the intra-subject variability of the behavioral study, for each of the *N*_simulations_ per pair, histograms of the parameters of the estimated psychometric functions from the computer simulation were compared to histograms of the parameters of the psychometric functions originating from the behavioral data. This was done by calculating the cosine similarity between the two vectors of histogram bin counts (**h** and **h**^*T*^, for the behavioral and simulated data, respectively) for the parameters *α, β*_inflections, b.c. [a.u],_ and *λ* :

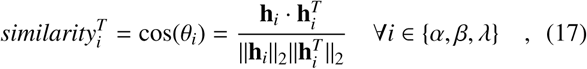

where a similarity of 1 represents identical histograms. Note that by using this similarity metric the histograms do not need to be additionally normalized. The following bin sizes were used for *α, β*_inflection, b.c. [a.u.]_, and *λ*: 0.25, 0.05, and 0.005. To obtain an overall similarity, the three calculated similarities were multiplied with each other.

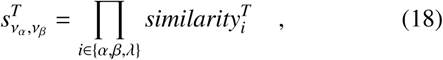

From these *N*_simulations_ overall similarity values, the arithmetic mean 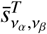 was calculated. The iso-reliability contour was projected onto the similarity surface in the two-dimensional noise space after a spline interpolation, identical to what was done for the reliability. The interpolated *ν*_*α*_ and *ν*_*β*_ pair on the iso-reliability contour with the highest average overall similarity was chosen as the best model to estimate intra-subject variability (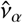 and 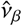).

For a new set of psychometric functions 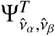 with the optimal noise model, the simulation was repeated *N*_simulations_ times, and the parameter distributions as well as 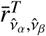 and 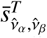 were calculated. In addition, the maximum attainable reliability 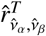 (corresponding to no method variability) was computed based directly on the templates with intra-subject noise, but without simulating the psychophysical experiment.

To illustrate the intra-subject variability on a psychometric function, a population average model was computed by averaging the individual subject models. Using the intra-subject variability models with parameters 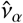 and 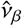, 1000 templates were created. The estimate distributions originating from pure method variability as well as from intra-subject variability were compared with each other by plotting the percentage of estimates within a tolerance interval depending on the interval size (percentage within bounds, *PCTw/iB*), and the normalized area under these curves (*nAUC*) according to the methods proposed by Rinderknecht et al. (in preparation).

## 3 Results

The test-retest reliability coefficient of the behavioral study and its confidence interval was *r* = 0.212 [0.077, 0.394]. The simulated reliability 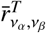 for different *ν*_*α*_ and *ν*_*β*_ pairs as well as the matched iso-reliability contour at *r* are shown in **Figure 2**. In case of no intra-subject variability, the reliability of the parameters of the psychometric functions originating would correspond to 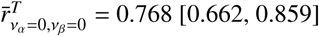 for the psychophysical paradigm and sampling procedure used in this work (i.e., maximum attainable reliability using these methods for the present population of interest). The overall similarity 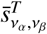 (combined for threshold, slope, and lapse rate) is visualized in **Figure 3**, together with the same projected iso-reliability contour. The maximum overall similarity on the contour was found for the noise level pair 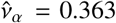 and 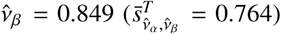, corresponding to the best intra-subject variability model estimate. The similarities of the distributions of the parameters of the psychometric functions are shown individually in **Figure 4**. The simulated reliability at this noise level pair was 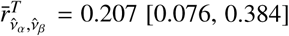. The maximum attainable reliability without method variability (i.e., assuming a perfect assessment) for the identified intra-subject variability model would be 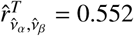. For illustration purposes, the effect of intra subject variability on the shape of the psychometric function is shown in **Figure 5** for the population average model *ψ*(*x*; *α* = 1.696, *β* = 1.708, *γ* = 0.500, *λ* = 0.036) and the noise level pair 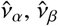, together with the distributions of threshold and slope resulting from method and intra-subject variability. For the threshold estimation, the *nAUC* was higher for the method variability compared to the intra-subject variability, whereas for the slope estimation, the opposite was the case. The maximum difference in estimation performance in terms of *PCTw/iB* was 42.5% at a threshold tolerance of ±0.210°, and 38.1% at a slope tolerance of ±0.299.

**Figure 2.**
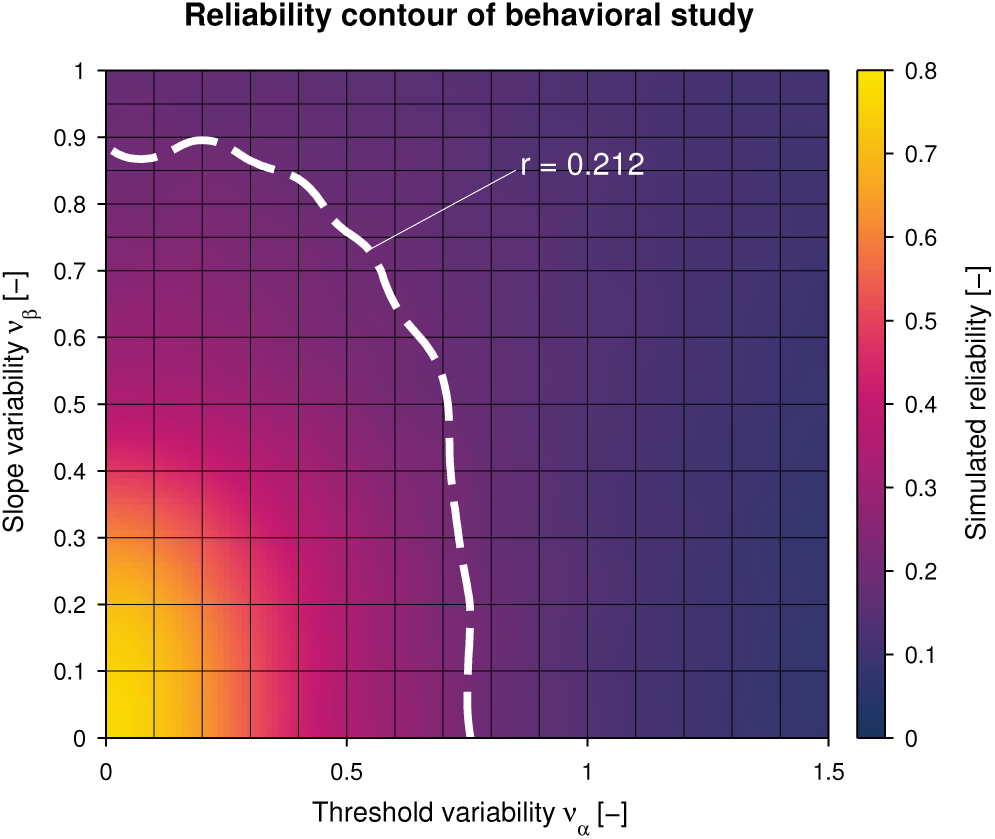
Simulated reliability and iso-reliability contour of behavioral study. For each pair of intra-subject threshold noise *ν*_*α*_ and slope noise *ν*_*β*_, the simulated reliability averaged across *N*_simulations_ = 1000 simulations 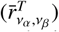 is represented as a heat map. The dashed white line indicates the iso-reliability contour corresponding to the reliability obtained from the behavioral study (*r* = 0.212).

**Figure 3.**
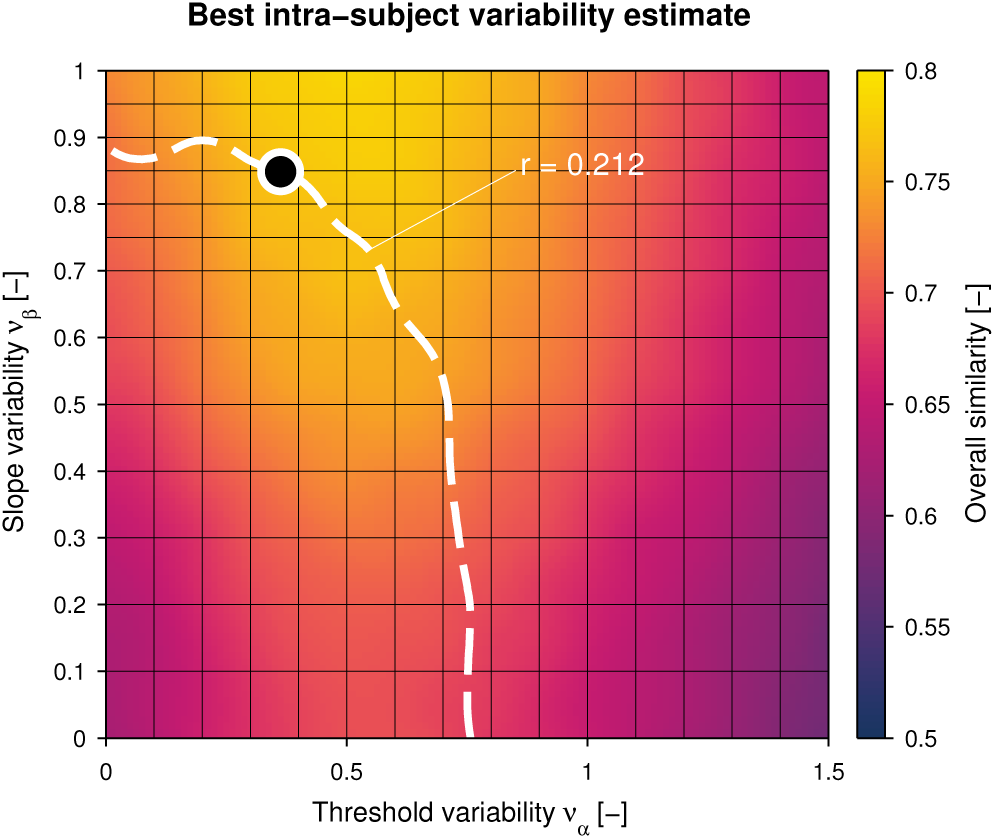
Best intra-subject variability estimate based on overall similarity. For each pair of intra-subject threshold noise *ν*_*α*_ and slope noise *ν*_*β*_, the overall similarity (combined for threshold, slope, and lapse rate) averaged across *N*_simulations_ = 1000 simulations 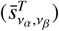 is represented as a heat map. The dashed white iso-reliability contour is identical to **Figure 2**. The noise level pair on the contour with the highest overall similarity 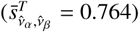 is indicated with a black dot.

**Figure 4.**
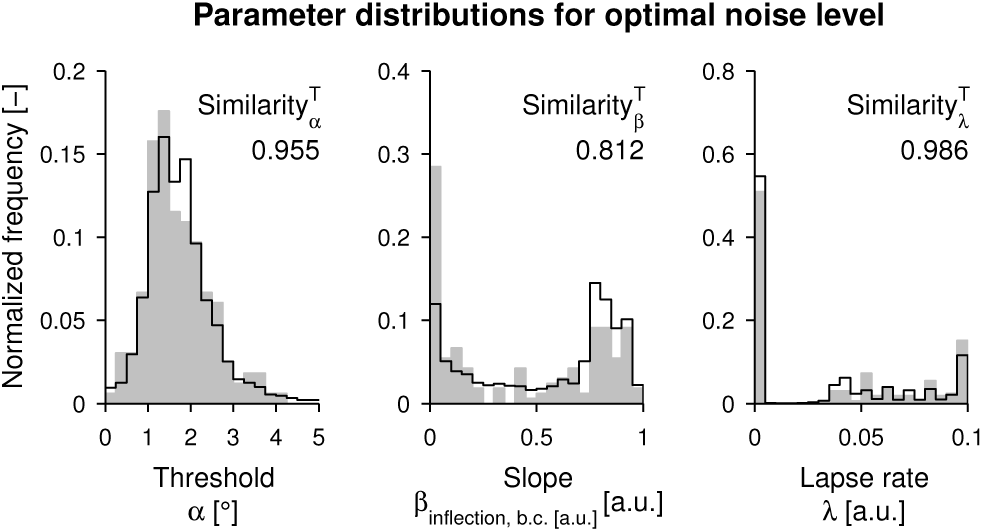
Histogram similarity for the optimal intra-subject variability. Histograms of the parameters of the psychometric functions of the behavioral data (gray fill, 33 × 5 data points) versus the simulated data (black outline, 33 × 5 simulated data points averaged over 1000 simulations) with optimal noise level at the pair 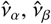

**Figure 5.**
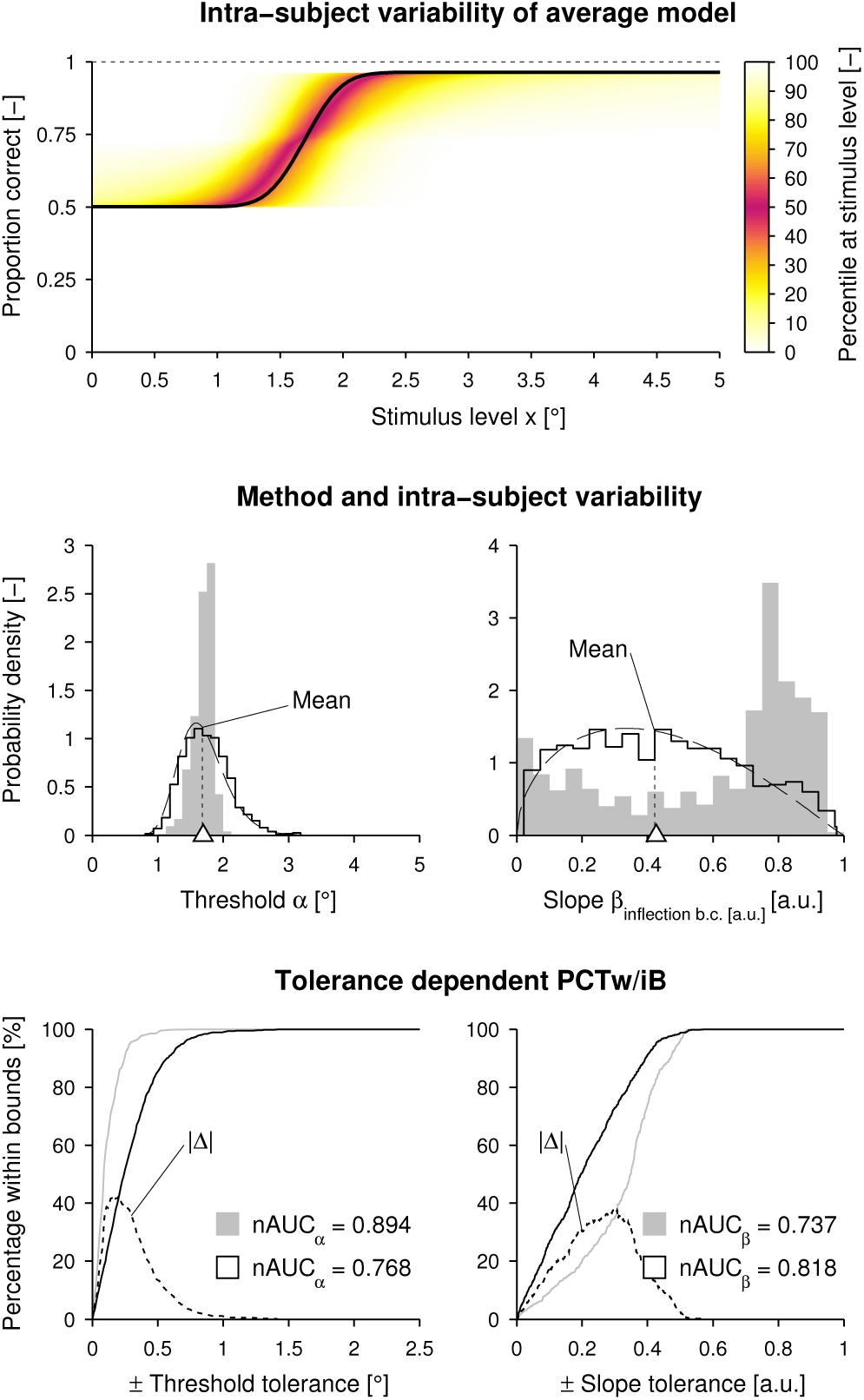
Illustration of intra-subject variability. (Top) For each stimulus level *x* the distribution of proportion correct is plotted as a heat map based on 1000 templates created with the population average model (bold black sigmoid) and the intra-subject variability models using 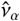 and 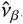. **(Middle)** The dashed black distribution curves correspond to the parametric log–normal and beta intra-subject variability models. The histograms (black outline) as well as the dashed black lines for the means show the parameter distributions of the 1000 templates including intra-subject variability. The white triangles indicate the threshold and slope of the population average model (without noise). As a comparison, the inherent method variability (histogram with gray fill) for the same population average model without intra-subject variability is plotted. (**Bottom**) The percentage of estimates within a tolerance interval (percentage within bounds, *PCTw*/*i*B) around the parameters of the population average model is plotted against the size of the interval (gray: method variability, black: intra-subject variability), together with the absolute difference of percentage (|Δ|, dashed black line). For both method and intra-subject variability, the normalized area under the curve (*nAUC*) is calculated.

## 4 Discussion

In this work we presented an approach to quantify intra-subject variability in psychophysical testing. This was achieved by introducing and adjusting a statistical noise model in computer simulations to match the test-retest reliability and histograms of the parameters of the estimated psychometric functions of a behavioral dataset. Using this approach we estimated the intra-subject variability of healthy subjects in a psychophysical assessment of proprioceptive perception at the wrist using a 2AFC paradigm, and compared the intra-subject variability with the inherent method variability of PEST.

The results showed that for a matched reliability, the similarity between the behavioral and simulated datasets was excellent for the optimal pair of intra-subject threshold and slope variability. Furthermore, the identified intra-subject variability of the threshold was larger compared to the method variability, whereas the opposite was the case for the slope.

### 4.2 Intra-subject and method variability

When trying to estimate the test-retest reliability based on the population model without intra-subject variability, the [0.403, 0.704]. For illustration purposes, the effect of intrareliability coefficient would be largely overestimated. In the present sample population this would result in a considerable error of 262.3%. In contrast, when including intra-subject variability in the simulation, the reliability of the simulated experiment matched the reliability of the behavioral study with an absolute error of 0.005 (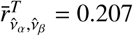 and *r* = 0.212, respectively), corresponding to a relative error of 2.4%. In theory, this error should be zero, however, since the estimates were based on a stochastic generation of responses, the simulated test-retest reliability varied across simulation runs. To improve the estimate of intra-subject variability, and therefore the match of reliability values, a high number of repetitions (*N*_simulations_) were performed to obtain higher statistical power, and the grid of the simulated intra-subject variability levels in the two-dimensional reliability space was interpolated. This error could be further minimized by increasing the number of repetitions and the density of the simulation grid. Further indication for a good model estimation quality is provided by the fact that not only the simulated and behavioral reliability coefficient matched, but also matching errors for the CI were low (absolute [0.001, 0.010] and relative [1.3%, 2.5%] errors for the lower and upper bound). Moreover, cosine similarity between behavioral and simulated outcome measures was very high for all three parameters *α, β*, and *λ* (*>* 0.8), and thus demonstrates that the population’s inter- and intra-subject variability models accurately represent the actual population.

The presented method allows to discern between and compare intra-subject variability and method variability. When assuming invariant subjects (i.e., no intra-subject variability), the test-retest reliability for the threshold would be 39.2% higher compared to when the estimated intra-subject variability is included in the simulation, but a perfect method (i.e., no method variability) would be assumed. This is also reflected by the *nAUC* for the threshold (a non-parametric metric to evaluate the variability of estimation errors), which is higher by 16.4% for the simulated case with method variability only. Based on these findings, if the assessment was to be improved, one could suggest to address factors influencing the intra-subject variability, before optimizing the psychophysical sampling procedure, as even with a perfect method, the reliability would ceil at 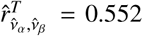 due to intra-subject variability. On the contrary, the slope estimates suffer from poor method performance and, according to the U-shaped estimate distribution (histogram with gray fill in **Figure 5 (Middle, right)**), outcome measures are predominantly severely under- or overestimated. As a consequence, the *nAUC* for the simulated case with intra-subject variability only is 11% higher. Thus, if the slope estimation should be improved, it would be important to optimize the current sampling procedure or choose another sampling procedure (e.g., the Ψ method, designed to estimate both the threshold and the slope (Kontsevich and Tyler, 1999)).

### 4.2 Advantages and limitations of this method

The advantage of this method is that the test-retest reliability is affected by all terms of variability (interand intrasubject, and method variability). As a consequence, since the inter-subject variability can be approximated by taking the averaged psychometric functions for each subject and the method variability is given by the simulation, the intra-subject variability can be estimated. Furthermore, the reliability can be calculated even if only two measurements were done per subject, whereas, for example, calculating the standard deviation of two measurements for each subject is very likely a poor estimate of the real variability (besides being still con-found with method variability). However, it should be noted that, depending on the intra-subject and method variability, the quality of the model of the population (and inter-subject variability) can be compromised if only two measurements are available per subject. Thus, in case of a poor population model, an overestimated inter-subject variability may be compensated by an underestimated intra-subject variability and vice versa when matching the reliability. Ideally, the available behavioral data would encompass a large sample size (for a good representation of the population) and a large number of measurements (for a good estimate of each subject’s psychophysical function). An advantage of sampling templates from the computed distributions representing the population compared to using the averaged psychometric functions as templates, is that repeated randomly sampled templates should lead to more generalizable results than bootstrapping from a limited set of subjects. More importantly, it offers the possibility to sample more templates from the distribution, for example to predict how the reliability and its confidence interval changes with increasing sample size.

A limitation of the present simulations is that no intra-subject variability was modeled for the lapse rate. It would be possible model the lapse rate including intra-subject variability with a beta distribution as for the slope, but with an adapted support. However, for the sake of simplicity, this was omitted here. As a matter of fact, as the histogram similarity of the lapse rate parameter is almost 1, it shows that adding a lapse rate variability model may not be necessary. When identifying the best model of intra-subject variability, the noise level pair *ν*_*α*_, *ν*_*β*_, where overall similarity is the highest, may not lie on the iso-reliability contour corresponding to the reliability *r* of the behavioral data. One reason is that in the histogram, similarity inter- and intra-subject variability are confounded, and the similarity may vary depending on the selection of bin sizes. In contrast, using the reliability as a metric should provide a more robust and accurate estimate of the variability model, as it distinguishes between inter- and intra-subject variability despite taking both into account. Therefore, the overall similarity is used only as a second criterion to find the optimal model. One major limitation of this approach to estimate intra-subject variability, is that it only provides one variability model for the whole sample and not individual models for each subject. However, this is already a significant improvement over no variability model, and may be accurate enough for many applications.

## 5 Conclusions

Computer simulations offer a valuable and powerful tool to simulate and optimize psychophysical experiments. While they can be used to evaluate different procedures and their method variability, existing computer simulations are often not representative of real-world scenarios, as critical aspects such as the intra-subject variability are neglected. As a matter of fact, intra-subject variability cannot be directly quantified from behavioral data. This work introduces a new approach based on the combination of computer simulations and behavioral data to separate method variability from intra-subject variability and to estimate and model intra-subject variability in psychophysical experiments.

Given a realistic model of the population, different psychophysical procedures can be simulated and compared, and the procedures can be tuned to the specific application and target population. Quantifying the method and intra-subject variability allows putting them into perspective when developing assessments. Given the intra-subject variability, it allows simulating an experiment with an ideal psychophysical method (i.e., finding the theoretically maximally attainable performance of an assessment). These two aspects can inform the decision whether effort should be spent on improving the psychophysical procedure (i.e., reducing method variability) or if potential confounds affecting intra-subject variability should be addressed. The efficiency of attempts to reduce confounds (e.g., inattention (Rinderknecht et al., 2018)) could be quantified (using the presented method) based on a reduction of the intra-subject variability. Furthermore, based on the more complete model also containing intra-subject variability, it is also possible to examine the impact of a larger number of trials on reliability, or the converging behavior of the reliability’s confidence interval bounds with a larger number of subjects, as well as retests, without having to conduct additional experiments. This presents a particular benefit for studies with populations where time for assessments is limited or expensive, as in the case of a clinical setting.

## Author contributions

MR, OL, and RG contributed to the conception of this work. MR developed the methodology, implemented the experiments and the computer simulations, performed the analysis, interpreted the results, and drafted the manuscript. MR, OL, and RG revised the manuscript and approved the final version.

## Acknowledgments

The authors would like to thank J. Egloff and S. Huber for the acquisition of the behavioral data as well as W. L. Popp for fruitful discussions. This research was supported by the ETH Zurich Foundation in collaboration with Hocoma AG, the Janggen-Pöhn Foundation, and the Swiss National Science Foundation through project 320030L_170163.

